# A fast and simple approach to *k*-mer decomposition

**DOI:** 10.1101/2024.07.26.605312

**Authors:** Patrick Kunzmann

## Abstract

Alignment searches are fast heuristic methods to identify similar regions between two sequences. This group of algorithms is ubiquitously used in a myriad of software to find homologous sequences or to map sequence reads to genomes. Often the first step in alignment searches is *k*-mer decomposition: listing all overlapping subsequences of length *k*. This article presents a simple integer representation of *k*-mers and shows how a sequence can be quickly decomposed into *k*-mers in constant time with respect to *k*.

## I. Introduction

A common task in bioinformatics is finding similar regions between sequences. In one scenario finding homologs of a sequence in another genome may reveal common functionalities between them. In another scenario the reads obtained from sequencing need to be mapped to a position on a reference genome to assemble a genome or quantify the number of transcripts. A *similar region* can be formalized as so called *alignment*: It specifies which position in one sequence corresponds to a position in the other sequence. The dynamic programming algorithm to obtain the guaranteed best alignment solution [1] is not computationally feasible for most modern applications: the length and number of sequences is simply too large.

To solve this problem, heuristic approaches emerged. Many modern algorithms (see [2], [3] as examples) build upon the concept of finding exact matches of length *k* between the sequences [4]. These subsequences of length *k* are termed *k*-mers. The process of finding all overlapping *k*-mers in a sequence is commonly titled *k-mer decomposition*.

**Example:**

Take the sequences `TATGC` and `ATGG`: Decomposition into 3-mers yields the enumeration^1^ (TAT, ATG, TGC) or (`ATG, TGG`), respectively. When comparing the enumerations, we find a match in the common `ATG` sequence.

To accelerate the identification of *k*-mer matches, the obtained *k*-mers are often stored in a *hash table* for fast lookup of the positions where a given *k*-mer appears. This requires *k*-mer *hashing*, i.e. mapping a *k*-mer to an integer, the *hash*. If this mapping is unambiguous, i.e. two different *k*-mers are guaranteed to get different integers assigned, it is called *perfect hashing*. If for *n* possible different *k*-mers the hash values are in the range 0 to *n* − 1, the hash is called *minimal perfect hash*. This is a favorable property, as it allows to implement the *hash table* as array of size *n*.

Although *k*-mer decomposition is the fundament of many modern alignment tools, most literature about their underlying algorithms focus on how the *k*-mers are used to find the alignment. This article puts the spotlight on *k*-mer decomposition itself: It presents an intuitive integer representation of a *k*-mer, which at the same time acts as minimal perfect hash. This is accompanied by a *minimal perfect hash function* (MPHF) that decomposes a sequence into these hash values in constant time with respect to *k*. Finally this paper proposes a simple way to give these *k*-mer hashes a pseudorandom ordering, a desirable property for certain *k*-mer based methods, such as *minimizers* [5] and *syncmers* [6].

The techniques presented in this article are also implemented in the *Python* bioinformatics library *Biotite* [7].

## II. Algorithm

### A. Sequence representation

Each sequence is subject to an *alphabet* Ω, that enumerates the symbols that are allowed in a type of sequence. Let the *symbol code* be the 0-based position of a symbol in Ω. Let the *sequence code* be the symbol codes for all symbols in the sequence.

**Example:**

Take the DNA sequence *S* = `TATGC`. The underlying alphabet comprises the nucleotides, hence Ω = (`A, C, G, T`). The symbol code *c* for the first symbol in the sequence *s* = `T`, is the 0-based position of it in Ω, hence *c* = `3`. Doing this encoding for all symbols in *S* yields the sequence code *C* = (`3, 0, 3, 2, 1`).

Specifically, in case that the sequence is represented as ASCII text ^2^, mapping a sequence *S* to sequence code can be implemented using fast array access as described in Fig. 1, in contrast to employing a slower associative array ^3^.

**Fig. 1:**
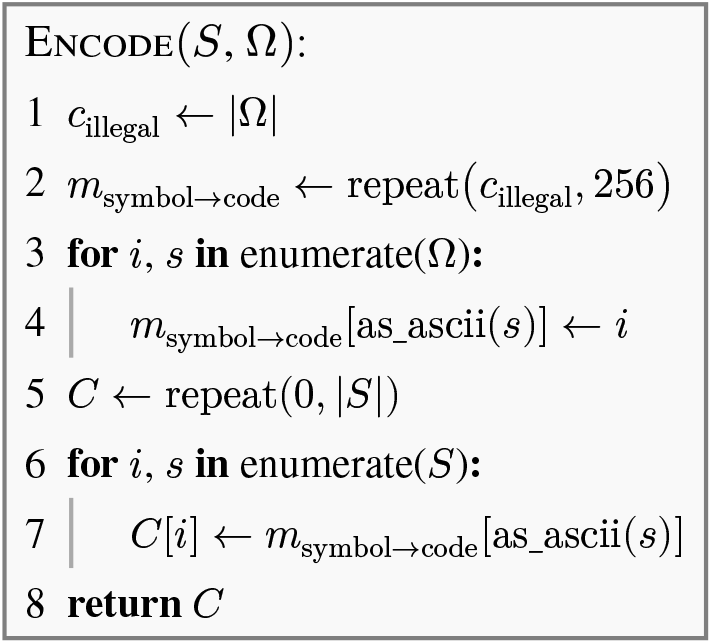
Sequence encoding into sequence code. The input sequence *S* is subject to alphabet Ω, which contains only ASCII-encoded symbols. As symbol codes are 0-based, the greatest symbol code is |Ω| − 1. Hence, *c*_illegal_ ← |Ω| can be used as marker value to check for symbols that are not part of Ω. The array *m*_symbol→code_ can be indexed with the ASCII code of a symbol to obtain the corresponding symbol code. For symbols that are not part of Ω, *m*_symbol→code_ would give *c*_illegal_.

### B. k-mer representation

The aim of the method presented in this article is to represent each *k*-mer unambiguously as single integer, that can be used as hash value. Analogous to the symbol code, this integer will be called *k-mer code*. First, the *k*-mer is converted into its sequence code as explained above. Next, the length of Ω, denoted by |Ω|, is used as radix to compute the *k*-mer code *c*^[k]^ from the sequence code *C* via

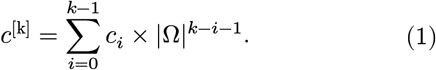

**Example:**

Take the 3-mer ATG that uses again Ω = (`A, C, G, T`) as base alphabet: The sequence code of this *k*-mer is (`0, 3, 2`). The corresponding *k*-mer code calculates as *c*^[k]^ = 0 × 4^2^ + 3 × 4^1^ + 2 × 4^0^ = 14.

Note that *c*^[k]^ can be again envisioned as element of an alphabet Ω^[k]^ that enumerates all possible *k*-mers. As Ω^[k]^ contains every combination of |Ω| symbols in each of its *k* positions, the length of such alphabet is

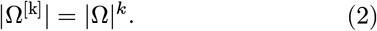

*C* can also be restored from the *k*-mer code *c*^[k]^ via

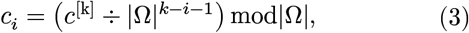

where ‘÷’ denotes integer division.

### C. k-mer decomposition

Performing *k*-mer decomposition of a sequence into *k*-mer codes requires application of (1) for each overlapping *k*-mer. Thus,

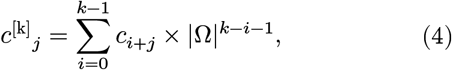

where *j* defines the 0-based sequence position where the *k*-mer starts. A naive implementation of this formula has a time complexity of *O*(*nk*), where *n* is the length of the sequence. However, it ignores the relation between two consecutive *k*-mer codes:

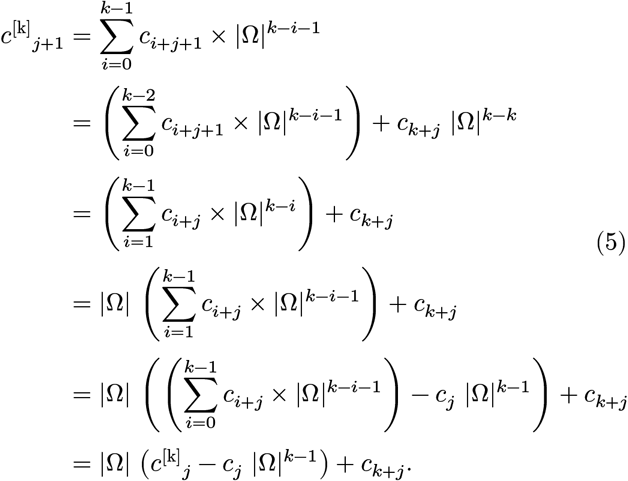

Intuitively, the *k*-mer code of the previous *k*-mer is taken, the symbol code of its first symbol is removed, the remainder is shifted to the left and the symbol code of the entering symbol is added.

As (5) contains no sum anymore, the time complexity is reduced to *O*(*n*). Instead the next code *c*^[k]^_*j*+1_ is computed from the previous code *c*^[k]^_*j*_. Only *c*^[k]^_0_ needs to be computed according to (4). In the implementation of (5) potentially further speedup can be achieved if |Ω| is a power of two. This is true e.g. for unambiguous nucleotide sequences with |Ω| = 4. In this case the compiler may substitute this multiplication with a fast bit shift operation depending on the hardware architecture.

Note that (5) only works for linear *k*-mers. Some algorithms in homology search use *spaced k*-mers [8], which contains ignored positions. In this case (4) can be still used.

### D. Pseudorandom ordering

In some scenarios the order of *k*-mer codes is relevant. For example minimizers [5] select only the smallest *k*-mer from a window of *k*-mers. This decreases the number of considered *k*-mers and in consequence improves the speed of finding *k*-mer matches between sequences. However, using the *k*-mer code directly to determine the ordering is not ideal: Especially, if the symbols in Ω are ordered alphabetically, the order in Ω^[k]^ is lexicographic. This means that *k*-mers with low complexity such as AAA… would always be the smallest *k*-mer, leading to more spurious than significant *k*-mer matches downstream [5]. A simple way to remedy this behavior is to apply a pseudorandom ordering to the *k*-mers, which can for example be achieved by choosing an appropriate *k*-mer hashing function [9].

However, in the presented case a *k*-mer is already represented as integer. Therefore, only an injective^4^ function *σ*(*c*^[k]^) is required to obtain an integer defining the ordering for a *k*-mer *c*^[k]^, i.e. the *k*-mer code *p* is defined to be smaller than *q* if *σ*(*p*) < *σ*(*q*). Furthermore, for the use case of computing sequence alignments, the quality of randomness is arguably less important than the speed of computation. Hence, a *linear congruential generator* (LCG) is appealing in this case. It predicts the next random number in a sequence via the simple formula

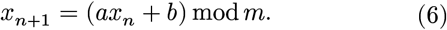

In a LGC with *full period* the sequence does only repeat after *m* elements. To achieve the full period attention has to be paid to the choice of *a* and *m*. Furthermore, *b* and *M* need to be coprime, which can be trivially achieved by setting *b* = 1 [10].

For the purpose of *k*-mer ordering, the LCG should be utilized to map a *k*-mer code *c*^[k]^ to a unique pseudorandom value *σ*(*c*^[k]^) that defines the *k*-mer ordering. Thus one can employ (6) to define

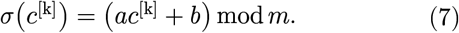

*σ*(*c*^[k]^) is only injective, if each *c*^[k]^ is mapped to a unique value. To ensure this property, an LGC with full period is used. If one handles up to 2^64^ *k*-mer codes^5^ *m* = 2^64^ is sufficient. When carefully implemented, the modulo computation in (6) is free due to automatic bit truncation. For *a* one can resort to published multipliers [11] that ensure both, full periodicity and good randomness for the chosen *m*. As example the following combination fulfills the requirements:

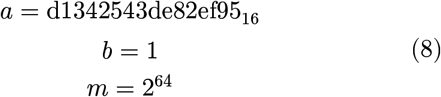

**Example:**

Take the nucleotide 3-mers `ATG` and `TGG` with corresponding *k*-mer codes *p* = 14 and *q* = 58, respectively. *σ*(*p*) = 8131822755183663655 and *σ*(*q*) = 7336488451890104259, with parameters taken from (8). This means *σ*(*p*) > *σ*(*q*). Thus, by the newly defined order, the 3-mer `TGG` is smaller than `ATG` in contrast to their lexicographic order.

## III. Results and Discussion

### A. Decomposition performance

The presented decomposition methods were benchmarked for different *k*-mer lengths on a 1000 bp nucleotide sequence as shown in Fig. 2. The benchmark was run on a system with *Apple M3 Max* processor @ 4.05 GHz using an implementation written in *Rust*^6^. As expected, the naive method scales linearly with *k* (*T* ≈ (1.00 + 0.38*k*) *μs, R*^2^ = 0.9997). In contrast, the fast decomposition method based on (5) runs in constant time (*T* ≈ 1.63 *μs*). As such potential optimization is hardware-related, this result may depend on the actual architecture and programming language.

**Fig. 2:**
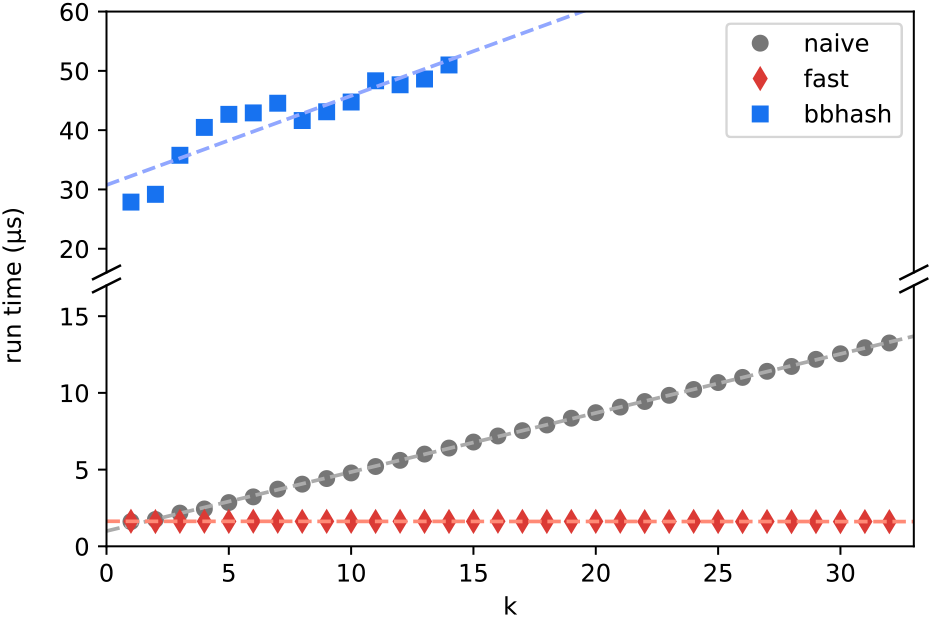
Run time of *k*-mer decomposition using different methods. Decomposition was run on a sequence with length 1000. The displayed run time includes also the conversion into sequence code. **naive**: Naive application of (4) for each sequence position. **fast**: Application of (5). **bbhash**: Application of *BBhash* [13], [14] on each *k*-mer string. The benchmark is limited to *k* ≤ 14 due to the memory requirements during construction of the hash function.

In summary, the fast decomposition method is already faster than the naive method for *k* ≥ 2, i.e. for any practically relevant *k*-mer length. The fast method is especially advantageous for algorithms that utilize long *k*-mers. For example, by default *Minimap 2* [15] uses *k* = 15 and *Kallisto* [2] uses *k* = 31. For these examples, the fast decomposition method is ∼4× and ∼8× faster than the naive method, respectively.

Note that the implementation used for the benchmark also includes sequence encoding itself: If the implementing library already stores sequences in their sequence code form, *k*-mer decomposition becomes faster than shown in the benchmark.

### B. The k-mer code as minimal perfect hash

In the framework of hashing the presented *k*-mer decomposition function (5) can be seen as MPHF, as long as only a single *k* is used:

- It is *perfect* as two different *k*-mers always get different *k*-mer codes.
- It is *minimal* as the *k*-mer codes range from 0 to |Ω^[k]^| − 1.

However, unlike other MPHFs (e.g. *BBhash* [13]), this MPHF produces hash values, i.e. *k*-mer codes, with the same lexico- graphic order as the input *k*-mers. Hashes with a pseudorandom ordering can be obtained by applying an LCG to the *k*-mer code according to (7). The resulting values are not minimal anymore though, as they are not within 0 and |Ω^[k]^| − 1, but they range between 0 and the LCG period *m* − 1.

This tradeoff can easily remedied by separation of concerns in the implementation: For building a *hash table* minimal perfect hashes are desirable, but random order is not required though. Hence, the original *k*-mer code can be used as keys here. On the other hand when requiring a random order, for example, to select the minimizer from *k*-mer codes [5], one can use (7) to obtain the order only. Downstream the original *k*-mer codes can be used again.

Apart from its simplicity, the advantage the *k*-mer decomposition method presented in this article over general purpose MPHFs such as *BBhash* is that it leverages the fact, that the objects to be hashed are simple enumerable *k*-mers. Hence, the construction of this MPHF is almost free. The only information required is Ω, which contains usually only a few symbols, and *k*. There is no costly construction time of the MPHF and its storage requirements are minimal, in contrast to general purpose MPHFs [16]. Furthermore, the more sophisticated computations of such MPHF also require more computation time: In the presented benchmark (Fig. 2), a *Rust* implementation of *BBhash* [14] required ∼20× longer than the fast method based on (5), even for *k* = 1 (*T* ≈ (30.75 + 1.50*k*) *μs, R*^2^ = 0.83).

## IV. Conclusion

This article advocates representing *k*-mers as integer in memory for mainly two reasons: First, using the presented algorithm *k*-mer decomposition can be achieved in constant time with respect to *k*. Since modern sequence alignment algorithms strive to be as fast as possible, this performance gain may pose a crucial advantage. Second, many current application of *k*-mers already at least implicitly rely on conversion of *k*-mers into an integer by means of hashing. Among other applications, this includes

- comparison of two *k*-mer sets to approximate sequence identity (e.g. [17]),
- efficiently finding match positions between two sequences using a *k*-mer lookup table (e.g. [3]) and
- map reads to genes using pseudoalignment (e.g. [2]).

Already having *k*-mers as unique integers at hand removes the need for hashing them and thus may further speeds up such applications.

In addition, representing a sequence as array of integers has the advantage of generalizing the definition of a sequence beyond simple text: If the alphabet Ω does contain other objects than single characters as symbols, e.g. arbitrary strings or integers, each enumeration of these objects can be considered a sequence. This allows alphabets to contain more symbols than the 95 printable ASCII characters, which would enable, for example, creating and representing more fine-grained structural alphabets [18]–[20] in the future.

## Footnotes

1 The more common mathematical term would be *‘sequence’*. For better distinction from the biological sequence definition, the term *‘enumeration’* is used in this article.

2 This is true for almost every sequence type encountered in biology.

3 This container type is also termed *map* or *dictionary* depending on the programming language.

4 one-to-one

5 |Ω|^*k*^quickly leads to an combinatorial explosion of |Ω^[*k*]^|, making 64-bit integers necessary.

6 The benchmark code as well the raw data is available at *https://github.com/padix-key/kmer-article* and as archive [12].

